# Sample size critically shapes the reliability of EEG case-control findings in psychiatry

**DOI:** 10.1101/2025.11.10.687610

**Authors:** Aida Ebadi, Sara Kaderi, Borja Rodríguez-Herreros, Nadia Chabane, Sahar Allouch, Mahmoud Hassan

**Affiliations:** MINDIG, F-35000, Rennes, France; Electrical and Computer Engineering Department, Rafik Hariri University, Mechref, Lebanon; Service des Troubles du Spectre de l’Autisme et apparentés, Département de psychiatrie, Lausanne University Hospital (CHUV), Lausanne, Switzerland; School of Science and Engineering, Reykjavik University, Reykjavik, Iceland

## Abstract

Electroencephalography (EEG) studies in psychiatry have produced highly inconsistent findings, particularly in case-control designs. Small sample sizes are widely assumed to contribute to this variability, yet their impact on the reliability of EEG group comparisons has not been systematically quantified at scale. Here, we address this key gap using a large multisite resting-state EEG dataset comprising 2,874 participants aged 5-18 years, including individuals with attention-deficit/hyperactivity disorder, autism spectrum disorder, anxiety, learning disorders, and healthy controls. We extracted a comprehensive set of spectral, temporal, and complexity EEG features and performed repeated case-control comparisons across a wide range of sample sizes using extensive random subsampling. Results from small samples were unstable, with inflated and highly variable effect sizes across iterations. By contrast, larger samples produced consistent and reproducible findings, converging on uniformly small but robust effects. Statistical power rose steeply with sample size, whereas false positive rates remained relatively stable. These results demonstrate the central role of sample size in shaping EEG study outcomes and challenge the utility of conventional case-control approaches for biomarker discovery in psychiatry.

## 1 Introduction

Electroencephalography (EEG) is well-positioned for clinical translation in psychiatry: it is inexpensive, portable, and scalable, enabling deployment beyond research centers and into routine care, including low-resource settings. Despite sustained interest, EEG-derived markers for neurodevelopmental and psychiatric conditions have not proven robust, replicable, or clinically actionable (Cortese et al., 2023; Yun, 2024). The dominant paradigm contrasts group averages, most often spectral power, between patients and healthy controls (HC), advancing statistically significant differences as candidate biomarkers. Yet replication is uncommon, reported effects vary in sign and magnitude across studies, and generalizability is limited. These inconsistencies are evident in well-studied cases. In attention-deficit/hyperactivity disorder (ADHD), many reports describe elevated theta and reduced beta power (often summarized as an increased theta-beta ratio), yet several studies fail to replicate this pattern or even report opposite trends (Arns et al., 2013; Poil et al., 2014). Similarly, in autism spectrum disorder (ASD), a proposed “U-shaped” spectral profile (higher delta, theta, and gamma power alongside reduced alpha) has been both supported and contradicted across studies (Dede et al., 2025; Kakuszi et al., 2024; Murray et al., 2025; Neo et al., 2023; Precenzano et al., 2020; van Diessen et al., 2015). These patterns point to a reliability problem rather than a simple discovery gap.

A primary driver of this issue is the widespread reliance on underpowered study designs in the context of small, heterogeneous effects. Typical case-control EEG-biomarker studies enroll around 30 participants per group (Clayson et al., 2019; Neo et al., 2023), a scale that severely constrains statistical power, thereby limiting reproducibility. Classical power analysis provides a framework for estimating the sample size required to detect a given effect at a specified significance level and desired power (Cohen, 2009). However, this approach assumes a known, fixed true effect size, an assumption that is difficult to meet in EEG research, where reported effect sizes are highly variable and often estimated from small, underpowered samples. While concerns about sample size and statistical power have been widely acknowledged in fields such as MRI and fMRI (Marek et al., 2022), a quantification of this problem is lacking for EEG case-control research.

This challenge is particularly pronounced in psychiatry, where substantial heterogeneity exists both within and across diagnostic categories. Individuals sharing a diagnosis can differ substantially in their underlying neurophysiology, and the same diagnostic label may correspond to multiple distinct profiles. Diagnostic boundaries are also porous: comorbidity between conditions such as ADHD, ASD, anxiety, and learning disorders is common, and overlapping symptom dimensions cut across categories. This heterogeneity and overlap complicate the search for disorder-specific EEG markers, since a single category may not map onto a coherent neural signature (Cuthbert & Insel, 2013; Mammarella et al., 2022).

Here, we address this methodological gap by systematically quantifying how sample size influences the reliability of case-control EEG studies. We leverage a harmonized, multisite cohort (n=2,874) comprising HCs and individuals with ADHD, ASD, anxiety disorders (ANX), and learning disorders (LD). We quantify how reliability changes as a function of group size by resampling the cohort across a wide range of *n* (from 10 to several hundred participants per group; 1,000 iterations) and re-estimating case-control effects. Analyses encompass spectral, temporal, complexity, and dynamical features. We evaluate significance rates as a function of sample size and sampling variability, and assess effect-size stability and inflation.

## 2 Methods

### 2.1 Dataset

The dataset analyzed in this study comprises 2874 individuals, including 558 HCs and 2316 patients diagnosed with ADHD (n=1221), ASD (n=554), ANX (n=305), or LD (n=236), aged between 5 and 18 (mean=10.00 ± 2.99, 42% M, Fig. 1a, Table S1). The cohort is compiled from five independent studies: the Healthy Brain Network Dataset (*<u>HBN</u>*) (Alexander et al., 2017; Langer et al., 2017), Multimodal Resource for Studying Information Processing in the Developing Brain (*<u>MIPDB</u>*) (Langer et al., 2017), Autism Biomarker Consortium for Clinical Trials Dataset (*<u>ABCCT</u>*) (McPartland et al., 2020), Multimodal Developmental Neurogenetics of Females with ASD (*<u>femaleASD</u>*) (Pelphrey, 2012), and *LausanneASD* dataset (Rodríguez-Herreros et al., 2025). Table S2 summarizes the distribution of data across contributing sites. For patients with multiple diagnoses (HBN, 41% of the cohort, Table S3, Fig. 1a), the diagnosis designated as DX_01 in the source dataset was retained. No participants were excluded on this basis for the main analysis. This dataset is the same as that used in our recent studies (Ebadi et al., 2025; Tabbal, Ebadi, Robert, et al., 2025).

**Fig. 1.**
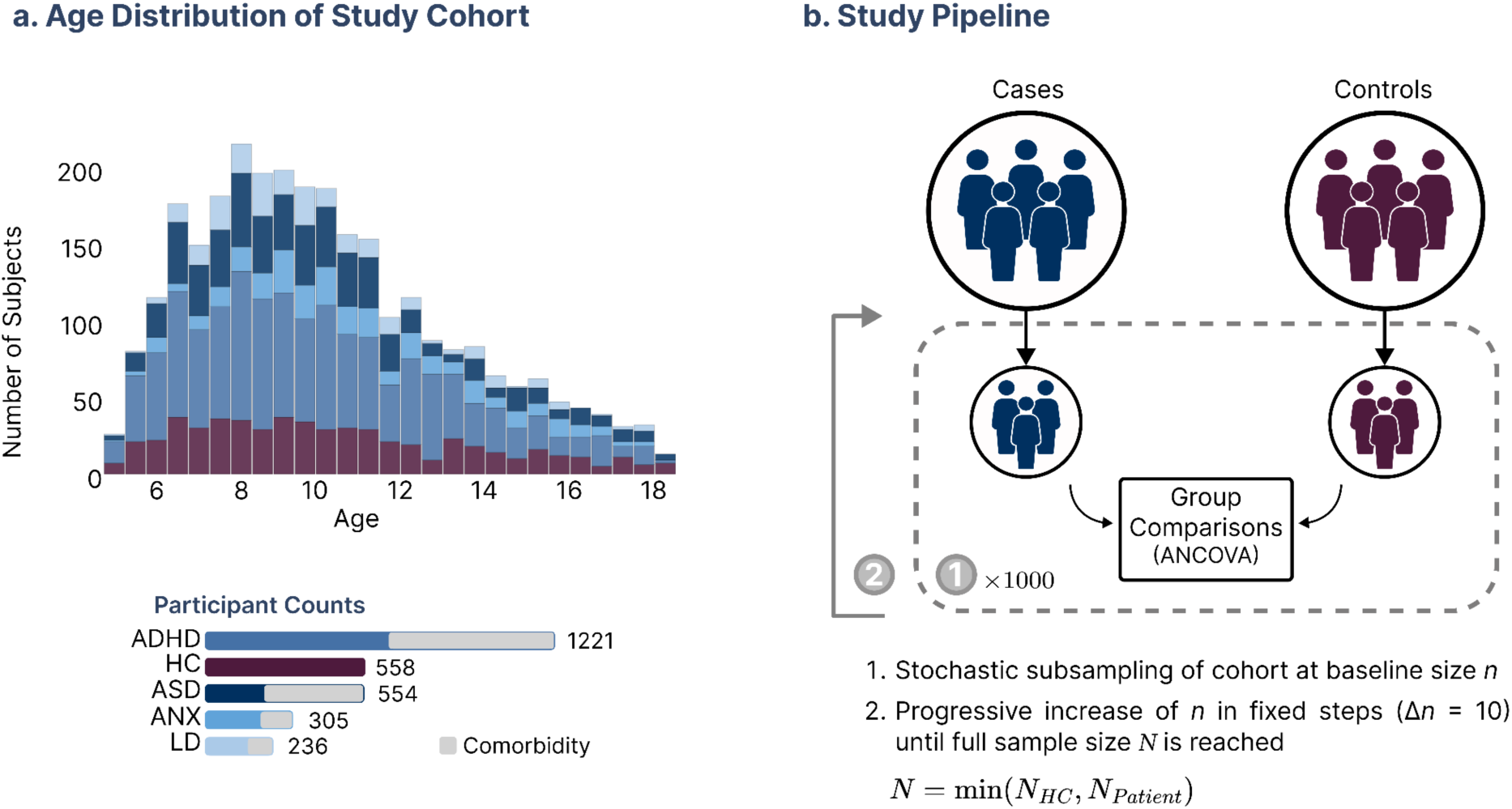
Study design and analysis pipeline. **a,** Demographic and cohort composition. Age distribution of participants and diagnostic group count, with consistent color mapping across panels. Comorbidity proportions are shown within each group. Exact sample sizes and demographic breakdowns are provided in Supplementary Table S3. **b,** Subsampling framework used to evaluate statistical stability across sample sizes. Cohorts were randomly subsampled at size *n*, repeated 1,000 times for each *n*, with sample size progressively increased in increments of Δ*n* = 10 until the full sample size *N* was reached. For each subsample, group differences were assessed using ANCOVA controlling for age and sex.

### 2.2 Preprocessing

Resting-state EEG was originally recorded with high-density 128-channel systems under eyes-open conditions and subsequently downsampled to 19 channels using the standard 10–20 montage. As source localization was not performed in this study, high spatial resolution was deemed unnecessary, and channel reduction allowed for substantial computational efficiency. The EEG preprocessing pipeline adhered to the automated approach described in our recent studies (Ebadi et al., 2025; Tabbal, Ebadi, Mheich, et al., 2025). Preprocessing steps included bandpass filtering (1–45 Hz), downsampling to 200 Hz, bad channel detection and interpolation using *pyprep* (Appelhoff et al., 2022; Bigdely-Shamlo et al., 2015), re-referencing to the common average, removal of ocular artefacts via Independent Component Analysis (ICA) with *IClabel* (Pion-Tonachini et al., 2019), segmentation into non-overlapping 10-second epochs, and rejection of artifactual epochs using *Autoreject* (Jas et al., 2017). A total of 175 participants were excluded after preprocessing due to either having more than 20% interpolated channels or exhibiting excessively noisy recordings upon visual inspection. Finally, a subject-wise z-transformation was applied: for each subject, the mean across all channels was subtracted and the result divided by the across-channel standard deviation, thereby normalizing EEG amplitudes, ensuring comparability.

### 2.3 Feature extraction

We extracted 103 EEG features spanning frequency, temporal, and complexity domains to comprehensively characterize the signals. Frequency-domain features included absolute and relative power computed across canonical frequency bands (delta [δ], 1–4 Hz; theta [θ], 4–8 Hz; alpha [α], 8–13 Hz; beta [β], 13–30 Hz; gamma [γ], 30–45 Hz), as well as adjusted power, alpha peak frequency, and several power ratios (θ/β, θ/α, δ/β, δ/α). We also included parameters of the aperiodic component (intercept and slope). Complexity features captured nonlinear and fractal properties of the signal. These included approximate entropy, sample entropy, spectral entropy, Hjorth parameters (activity, mobility, complexity), Hurst exponents (H, c), Katz’s and Higuchi’s fractal dimensions, detrended fluctuation analysis (DFA), and Lempel–Ziv complexity, each computed on broadband as well as frequency-specific signals (δ, θ, α, β, γ). Time-domain features included amplitude total power, mean and standard deviation of the envelope, and higher-order statistics such as skewness and kurtosis, all computed on broadband and frequency-specific signals. A substantial portion of the selected features was adapted from (Gordillo et al., 2023). In addition, we incorporated metrics from the CAnonical Time-series CHaracteristics (Catch22) toolbox (Lubba et al., 2019). This curated set of 22 features was selected from an initial pool of 4,791 descriptors for their broad discriminative power across classification tasks. Together with the signal mean and standard deviation, this yielded 24 time-series features. Full details of Catch22 features are available <u>online</u>. This feature set was motivated by three considerations: broad coverage across various domains; grounding in the established EEG literature, with the majority of features drawn from or inspired by prior EEG studies to ensure alignment with commonly reported measures (Gordillo et al., 2023); and the inclusion of Catch22 as an exploratory complement, together providing a comprehensive and representative characterization of resting-state brain activity.

Given the multi-site nature of our dataset, we applied feature harmonization using neuroCombat (Fortin et al., 2018; Jaramillo-Jimenez et al., 2024) to reduce site-related variability while preserving biologically relevant variance. Age was specified as a continuous covariate, while sex and group were included as categorical covariates to ensure that their effects were retained after harmonization. Results demonstrating the effectiveness of the harmonization procedure are provided in the Supplementary Materials (Fig. S1). We ultimately obtained 103 features per channel across 19 channels, along with their channel-averaged counterparts, for a total of 2,060 features.

### 2.4 Repeated subsampling framework

The proposed data-driven framework, inspired by (Marek et al., 2022), is illustrated in Fig. 1. In each iteration, 10 subjects were randomly drawn from the HC group and 10 from the patient group, followed by a group comparison using ANCOVA. This was repeated 1,000 times, with a new random sample drawn on each iteration. The *p-*value and effect size are recorded for each iteration. Subsequently, the subsample size is increased by increments of 10, and the procedure is repeated, up to the maximum number of subjects available in each condition. We performed ANCOVA using the Python *pingouin* library (Vallat, 2018). Age and sex were included as covariates to account for their effects when comparing the HC and patient groups. In the present study, the term ‘*group*’ refers to the diagnostic/status category of each participant.

Alongside *p-*values, we quantified effect sizes for each comparison. We used partial eta squared (η²ₚ) as the effect size metric, representing the proportion of variance in the outcome attributable to group differences, after accounting for covariates (Richardson, 2011). Values of η²ₚ are commonly interpreted as small (0.01–0.06), medium (0.06–0.14), and large (≥ 0.14) effects (Lakens, 2013).

We calculated a *significance percentage* for each EEG feature, defined as the proportion of 1000 resampling iterations in which a group comparison yielded *p* < 0.05. This metric quantifies the reproducibility of a given group difference across random subsamples of a given size, rather than relying on a single statistical test. A sample size threshold was defined as the smallest sample size at which ANCOVA yields a statistically significant difference in more than 95% of the iterations.

### 2.5 Inference error and spatial stability metrics

To comprehensively characterize the reliability of case-control EEG comparisons as a function of sample size, we quantified five metrics spanning statistical inference and spatial pattern stability.

**Statistical inference metrics.** We quantified three inference error metrics as a function of sample size: false negatives, false positives, and statistical power. Truly significant features were defined as those consistently reaching significance (≥95% of iterations) when tested at the full sample size. At each subsample size, false negatives were defined as the proportion of sampling iterations in which truly significant features failed to reach significance, while false positives were defined as the proportion of iterations in which non-significant features were incorrectly identified as significant. Statistical power was calculated as the complement of the false negative rate. Figures in the main text represent averages of these metrics across all features.

**Spatial stability metrics.** To characterize the spatial stability of group comparisons across resampling iterations, we computed two complementary metrics of channel-level reproducibility: Jaccard similarity and normalized entropy of the significant channel set. For each resampling iteration at a given subsample size, the significant channel set was defined as the set of channels for which ANCOVA yielded *p* < 0.05. The reference pattern was defined as the set of channels reaching significance in at least 95% of iterations at the full sample size. Jaccard similarity was computed between the significant channel set of each iteration and this reference pattern as J = |A ∩ B| / |A ∪ B|, where A is the significant channel set of a given iteration, and B is the reference pattern, yielding values between 0 (no overlap) and 1 (perfect agreement). Normalized Shannon entropy of the significant channel set was computed across iterations at each subsample size to quantify the diversity of spatial patterns produced by different subsamples, with 0 indicating identical patterns across all iterations and 1 indicating maximum diversity. Both metrics were averaged across features and reported as a function of sample size.

### 2.6 Effect size inflation

We quantified inflation by comparing effect sizes estimated at each subsample size to those obtained from the full sample. For features deemed significant, inflation was expressed as the ratio of the subsample effect size to the full-sample effect size. For each sample size, we recorded the proportion of iterations in which this ratio exceeded thresholds of 1.5, 2, or 5, and then averaged these proportions across all significant features. Because the maximum number of participants differed between patient groups and HCs, the full-sample effect size was defined as the mean across the final subsample iterations. Features considered significant at the full sample were those surpassing the 95% consistency threshold.

### 2.7 Comorbidity and epoch length

To assess the robustness of our findings, we performed two complementary control analyses.

**Comorbidity.** Comorbidity is pervasive in psychiatric populations and could, in principle, undermine the robustness of group comparisons by introducing additional heterogeneity and contributing to small or inconsistent effects. To assess this, we focused on the largest clinical subgroup in our dataset-participants diagnosed with ADHD from the HBN cohort. Using ANCOVA with age and sex as covariates, we compared monodiagnostic ADHD individuals with those with at least one additional psychiatric diagnosis. We then repeated the case-control analysis at full sample size, restricting the ADHD group to monodiagnostic participants and comparing them to HCs.

**Epoch length.** Epoch length is a known methodological factor that can significantly influence the estimation of EEG features (Levy, 1987). To investigate its effect, we recalculated EEG features using non-overlapping epochs of 2, 4, 6, and 8 seconds and compared their distributions to those obtained from our primary analysis, which employed 10-second epochs (Kolmogorov-Smirnov test with Benjamini-Hochberg false discovery rate correction). To determine the impact of these differences on study outcomes, we repeated the ANCOVA analyses for each epoch length and compared the results to the primary analysis. Analyses were conducted on the full sample with 500 resampling iterations.

All control analyses used the same ANCOVA implementation as the main analysis.

## 3 Results

### 3.1 Small sample sizes destabilize group comparisons. Consistency emerges only at larger sample sizes

To examine how sample size influences the stability of case-control EEG analyses, we applied a resampling framework in which subsamples of varying size (*n* = 10 to ∼550 per group, in steps of 10) were repeatedly drawn (1,000 times) and tested with ANCOVA (see Methods, Fig. 1b). Results from smaller sample sizes were highly variable and unreliable. Using relative alpha power (averaged across all channels) as an illustrative feature, we observed substantial fluctuation in both effect size and significance across the 1,000 subsamples. At *n* = 30 per group (reflecting the average sample size commonly used in EEG case-control studies), comparisons between HCs and individuals with ASD produced inconsistent outcomes: depending on the subsample, the ASD group appeared to have higher, lower, or statistically indistinguishable alpha power relative to controls. In contrast, analyses of the full dataset showed that HCs exhibited higher alpha power (Fig. 2a). In addition, the spatial pattern of group differences was also unstable. The specific scalp channels identified as significant varied considerably across subsamples and often failed to reflect the topographic profile observed in the full cohort (Fig. 2a).

**Fig. 2.**
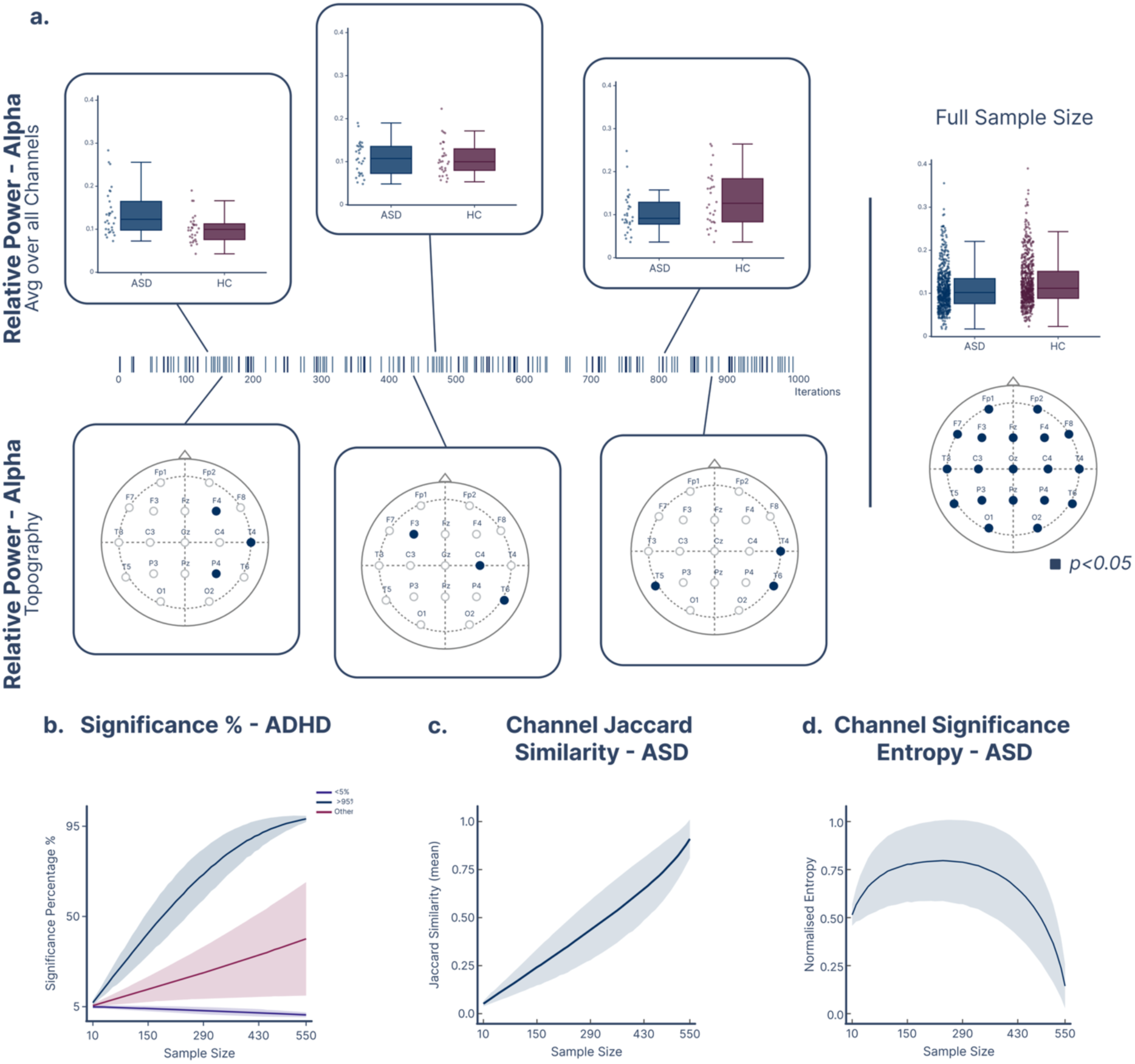
Overview of challenges associated with small sample sizes and summary of the dataset. **a,** Example feature (relative alpha power, averaged across all channels) illustrating instability across subsamples: boxplots from three example subsampling iterations at *n* = 30 per group show inconsistent effect directions and topographies, whereas the full dataset reveals a statistically significant difference (higher alpha power in HCs). Inset tick marks show a zoomed-in view of the significance pattern for an example feature (as in Fig. 3a–d), where each tick indicates a subsampling iteration that yielded *p* < 0.05. **b,** Significance percentage averaged across all features as a function of sample size for the ADHD group, illustrating how the proportion of iterations yielding *p* < 0.05 evolves with increasing n. Features are categorized into three profiles: those consistently reaching above 95% significance (dark blue), those remaining below 5% (purple), and values in between (red). Shaded areas indicate standard deviation across features within each category. **c,** Jaccard similarity between the significant channel set identified in each resampling iteration and the reference pattern derived from the full sample, averaged across all features for the ASD group. Values near zero at small *n* indicate that subsamples identify spatial patterns almost entirely different from the full-sample reference, while values approaching 1 at large *n* reflect convergence toward the true topographic pattern. **d,** Normalized entropy of the significant channel set across resampling iterations as a function of sample size for the ASD group. The inverted U-shaped profile reflects maximum spatial uncertainty at intermediate sample sizes, with entropy declining at larger *n* as a consistent topographic pattern emerges.

This pattern is reflected not only in the relative power of alpha but across the remaining features. Fig. 2b shows the proportion of significant findings across sample sizes, averaged over all features for ADHD; results for other diagnostic groups are shown in the Supplementary Materials (fig. S2). The significance pattern of 47% of the features differs markedly between lower and higher sample sizes, indicating that conclusions drawn from small samples are unlikely to generalize to larger cohorts. Furthermore, the spatial patterns emerging at lower sample sizes are highly inconsistent with those obtained at the full sample size.

Jaccard similarity, quantifying the overlap between the significant channel set identified in each resampling iteration and the reference pattern derived from the full sample, was near zero at small sample sizes, indicating that subsamples at small *n* identified channel sets almost entirely different from the true full-sample pattern (Fig. 2c). Similarity increased monotonically with sample size, approaching 0.9 at the maximum available *n*, consistent with progressive convergence toward a stable spatial pattern. Normalized entropy of the significant channel set showed an inverted U-shaped profile as a function of sample size (Fig. 2d). Entropy was low at small sample sizes, where few channels reached significance, peaked at intermediate sample sizes reflecting maximal spatial uncertainty, and declined at larger sample sizes as a stable topographic pattern emerged. Together, these results demonstrate that small sample sizes not only destabilize the statistical outcome of group comparisons but also produce highly variable and unreliable spatial patterns, undermining the interpretability of topographic findings reported in underpowered EEG studies. Similar trends were observed across the remaining diagnostic groups (Fig S3).

Having established that small samples yield unstable outcomes, we next quantified how statistical reproducibility changes with increasing sample size by calculating the *significance percentage* for each EEG feature. This allowed us to quantify how consistently each feature produced statistically significant differences as sample size increased. Across features, we identified four distinct reproducibility profiles (Fig. 3). Some features showed a monotonic increase in significance percentage with larger *n* (Fig. 3a,b), in some cases exceeding 95% (Fig. 3a). We define the *sample size threshold* as the smallest sample size at which this 95% level is crossed. In contrast, other features exhibited persistently low significance percentages across all *n*, indicating no significant group difference (Fig. 3c,d). A third pattern exhibited declining significance percentages with increasing sample size, suggesting initial false positives at low *n* (Fig. 3c). Finally, some features showed irregular, non-monotonic patterns, reflecting unstable group effects across subsampling (Fig. 3d). These profiles correspond to the three categories of features summarised in Fig. 2b: those surpassing the 95% threshold (Fig. 3a) to the upper line, those remaining near the 5% level (Fig. 3c) to the lower line, and the declining and irregular profiles (Fig. 3b,d) to the intermediate line. Their prevalence varied across diagnostic groups. In ADHD, 5.83% of features surpassed the 95% threshold, 52.82% remained below 5%, and 41.36% fell into the intermediate category; the corresponding proportions were 25.87%, 67.23%, and 6.89% in ASD; 7.43%, 58.45%, and 34.13% in anxiety; and 1.46%, 58.40%, and 40.15% in learning disorder. Notably, these reproducibility profiles did not appear to be specific to any disorder, feature type, or scalp location, prompting a more detailed analysis of the features and channels contributing to consistent effects. Next, we quantified the number of reproducible features per diagnosis and examined the sample size thresholds and spatial patterns associated with reliable group differences.

**Fig. 3.**
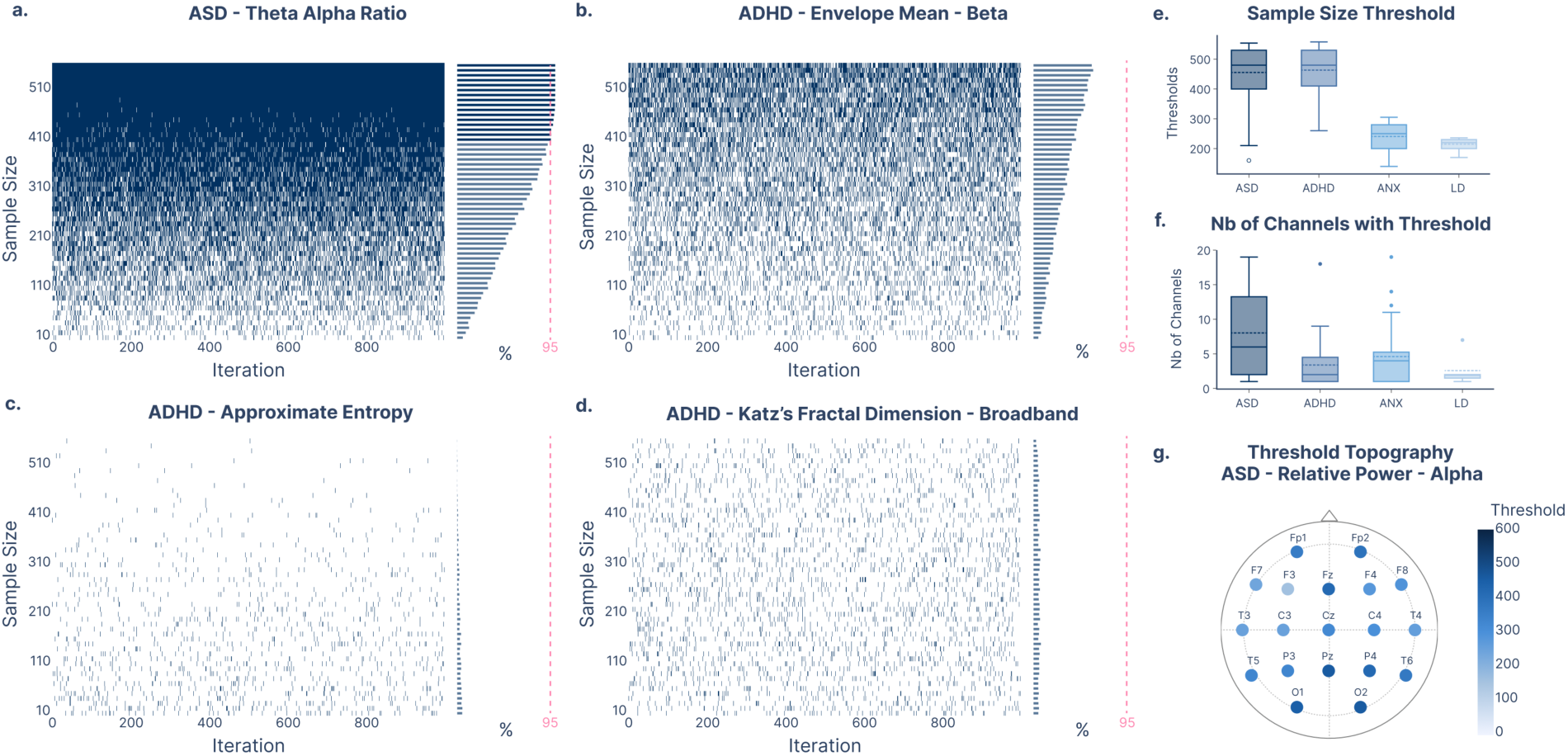
Overview of significance patterns. **a,** Examples of four reproducibility profiles, illustrating how significance rates evolve with increasing sample size across iterations (shown for **a,** ASD: theta–alpha ratio; ADHD: **b,** beta envelope mean, **c,** approximate entropy, and **d,** broadband Katz’s fractal dimension). **e,** Distribution of sample size thresholds for each diagnostic group, defined as the minimum *n* at which ≥95% of resampling iterations yielded p < 0.05. **f,** Distribution of the number of scalp channels per feature that reached 95% significance consistency, for each group. **g,** Topographic map of sample size thresholds for relative alpha power in ASD, showing spatial heterogeneity across channels.

Across the full sample, we identified a wide range of EEG features that showed significant group differences: 67 features for ASD, 37 for ADHD, 33 for ANX, and 12 for LD. For features with consistent group effects, we estimated the reproducibility threshold, the minimum sample size at which significance was achieved in 95% of resampling iterations. These thresholds were markedly higher than typical EEG study sizes: mean ± SD = 478.4 ± 48 for ASD, 490.1 ± 46 for ADHD, 255.7 ± 30 for ANX, and 217.6 ± 11 for LD (Fig. 3e). Spatial variability was also prominent: even within a single feature, different channels required different sample sizes to reach reproducibility (Fig. 3g). This heterogeneity was further reflected in the number of scalp channels contributing to significant effects. ASD showed the broadest spatial distribution, with a median of 8.0 significant channels per feature (Fig. 3f), whereas ADHD and ANX involved fewer channels, and LD was typically restricted to a single significant site.

Although several EEG features reached reproducibility thresholds within individual case-control comparisons, these effects were not diagnostically specific. Features showing consistent group differences were often shared across multiple diagnostic groups rather than being restricted to a single disorder (32.8% of the significant features), suggesting that the detected EEG alterations largely reflected transdiagnostic rather than disorder-specific effects.

### 3.2 Effect sizes at lower sample sizes are exaggerated

We systematically quantified false negatives, false positives, and statistical power across varying sample sizes. At *n* = 30, the average power for detecting significant EEG features was only ∼10%, while the false positive rate for ASD comparisons was 4.86%. As sample size increased, statistical power rose steeply, whereas the false positive rate increased only modestly. This pattern, rapid gains in sensitivity, limited inflation of false positives, and a sharp reduction in false negatives, was consistent across all diagnostic groups (Fig. 4).

**Fig. 4.**
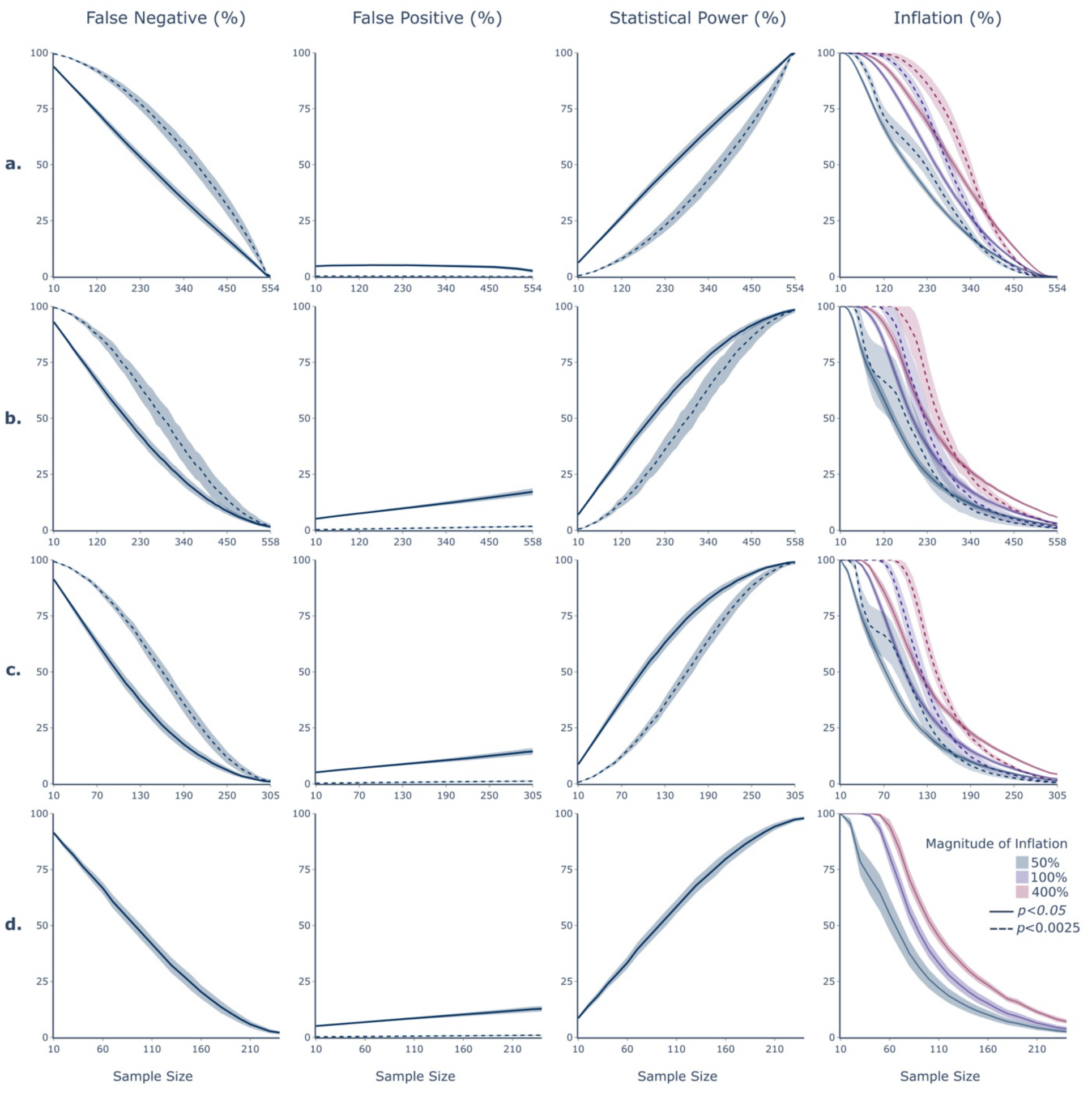
Inference error (false negatives, false positives, statistical power) and effect size inflation profiles across groups. Profiles are calculated across all significant features. **a,** ASD **b,** ADHD **c,** ANX **d,** LD

Beyond statistical significance, the magnitude of observed group differences is crucial for interpreting clinical and biological relevance. We therefore examined how sample size influences effect size estimates (η²ₚ). Across diagnostic groups and EEG features, effect sizes were consistently and substantially inflated at small sample sizes. At *n* = 30, 98.6% of significant features exhibited effect sizes more than five times larger than those observed in the full dataset (Fig. 4a, Inflation %). This inflation was also accompanied by high variability in effect size estimates across subsamples. At n = 30, all effect sizes fell within the medium range, with no overlap with the full-sample distribution, where values were uniformly small and never exceeded 0.06 (Fig. 5a). As sample size increased, effect sizes converged toward their true values, revealing an inverse relationship between effect magnitude and the minimum sample size required for stability: robust effects stabilized quickly (i.e, reached the 95% consistency threshold), whereas subtle differences required much larger cohorts (Fig. 5b).

**Fig. 5.**
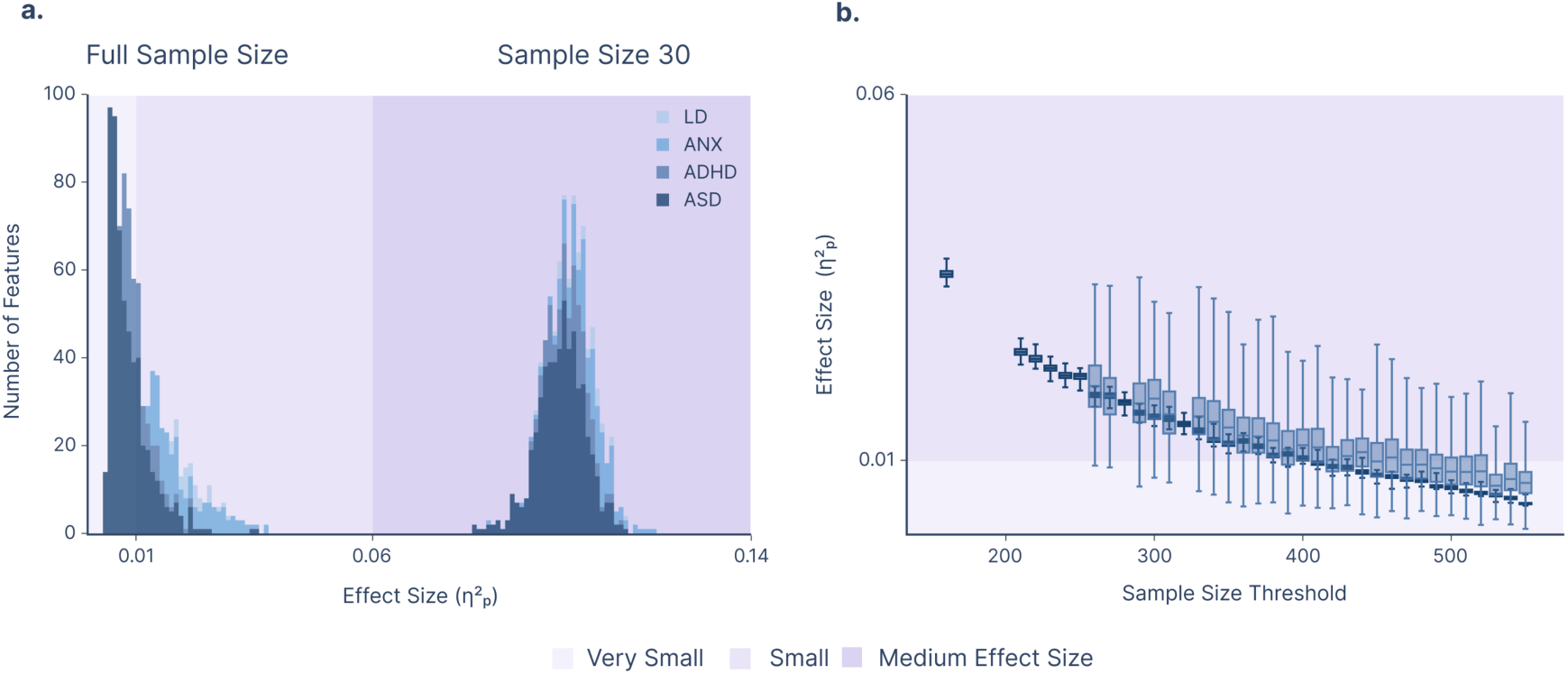
Effect size inflation and stability as a function of sample size. **a,** Histograms of effect sizes (η²ₚ) for features that were significant in the full sample, comparing the full dataset with analyses performed on subsamples of 30 participants per group. **b,** Distribution of full-sample effect sizes plotted against sample size thresholds for ASD and ADHD groups (for illustration, one representative feature is shown per sample size; all features exhibited the same trend).

### 3.3 Comorbidity has limited influence on case-control differences

Given the high prevalence of comorbidity in psychiatric populations, we next evaluated whether it contributed to the small or inconsistent group effects observed in our main analyses. Focusing on the ADHD subgroup, we compared EEG features between monodiagnostic individuals and those with additional psychiatric diagnoses, and examined how this distinction affected ADHD-control contrasts. Visual inspection of feature distributions revealed minimal differences between the groups, a pattern confirmed by ANCOVA controlling for age and sex. Only 114 out of 2,060 feature-channel pairs achieved significance (*p* < 0.05), all with negligible effect sizes. We then evaluated how comorbidity influences ADHD vs. HC comparisons. Restricting the ADHD group to monodiagnostic individuals altered the significance status of 197 features relative to the original analysis, with some previously significant features becoming non-significant and vice versa. However, these changes were not accompanied by meaningful changes in effect magnitude: nearly all features remained in the negligible range of effect size, with only one exceeding the threshold for small effect size. Thus, while comorbidity is highly prevalent and often invoked as a source of inconsistency in psychiatric research, its impact on EEG-based case-control differences in our dataset is limited. It is therefore unlikely to account for the small, unstable effects we observe.

### 3.4 Epoch length influences features but not group differences

Because epoch length can affect EEG feature estimation, we tested whether varying it (2–8 s, compared to the 10-s default) altered our findings. Feature distributions were indeed sensitive to epoch length: across 2,060 features, we found that 88.45%, 71.89%, 57.57%, and 40.48% of features differed significantly from the 10-second epoch features for 2, 4, 6, and 8-second epochs, respectively (Kolmogorov-Smirnov test with Benjamini-Hochberg false discovery rate correction). However, the downstream impact on case-control outcomes was limited: across all case-control comparisons (8240 comparisons including all of the four clinical groups), 5.6% of features changed in statistical significance on average (2s: 6.80%, 4s: 5.47%, 6s: 5.27%, 8s: 4.84%), that is, features that were previously significant became non-significant and vice versa. Crucially, these shifts were not accompanied by meaningful changes in effect size. As shown in Fig. 6, group differences consistently converged toward very small magnitudes, regardless of epoch length.

**Fig. 6.**
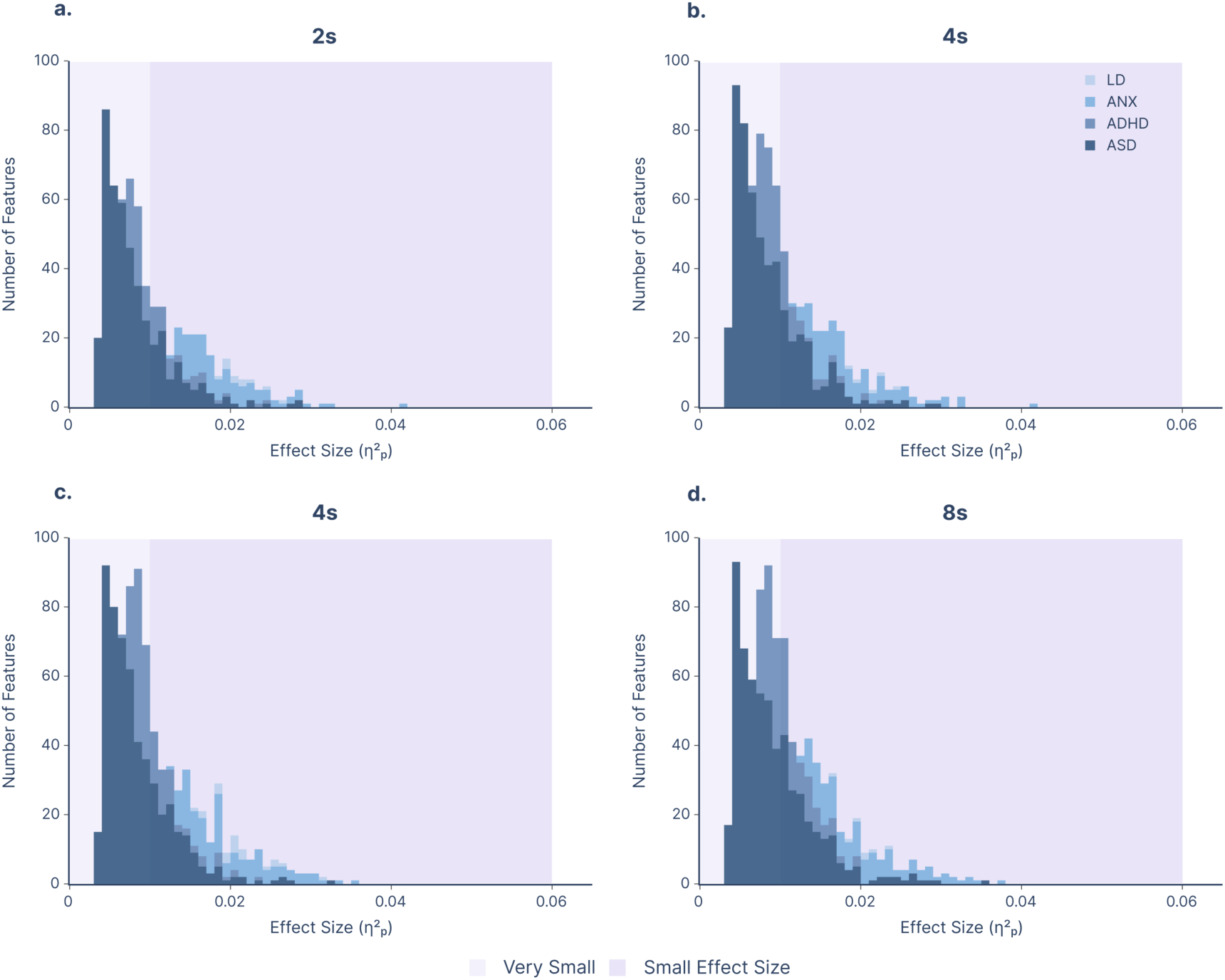
Histograms of significant effect sizes across different epoch lengths. Effect sizes (η²ₚ) are averaged over 500 resampling iterations at full sample size. Density plots are shown for the 103 features, averaged across all channels and min–max normalized for visualization. Panels illustrate results for **a,** 2s; **b,** 4s; **c,** 6s; and **d,** 8s epochs.

## 4 Discussion

In this study, we developed a sample-size-varying bootstrapping framework to systematically assess how sample size influences the reliability of resting-state EEG case-control comparisons across four neurodevelopmental and psychiatric conditions. At small sample sizes, case-control comparisons produced unstable results. In some cases, this instability persisted across all sample sizes tested; in others, increasing *n* led to more consistent outcomes. Across all features and diagnostic groups, the sample sizes required for 95% of subsampling iterations to yield significant group differences were substantially higher than the typical sample sizes used in EEG research, which often range from ∼20 to ∼30 participants per group (Clayson et al., 2019; Neo et al., 2023). While some group differences emerged at sample sizes as low as 140, the median sample size threshold for ASD and ADHD approached ∼480 participants, nearly the maximum available in our dataset for HCs (i.e., ∼550). Importantly, this threshold should not be interpreted as the minimum required number of subjects to detect a given effect for a specific feature. Rather, it underscores the magnitude of sampling error and the types of inferential pitfalls that arise at the sample sizes commonly employed in EEG research.

An important aspect of group comparison studies is the magnitude of the observed effects. Our results demonstrated that the effect sizes associated with significant findings were consistently inflated at lower sample sizes and exhibited considerable variability. This inflation is not random but systematically self-reinforcing: at small *n*, only those features with spuriously large estimated effects surpass the significance threshold, leading to a biased subset of results (Button et al., 2013; Marek et al., 2022). As sample size increased, the variability decreased, and estimates progressively stabilized, converging toward uniformly small values. These findings illustrate a key limitation of relying on statistical significance alone, especially as increasing sample size amplifies the likelihood of detecting effects that are statistically reliable yet practically negligible (Reddan et al., 2017; Sullivan & Feinn, 2012). In such contexts, *p-*values may be driven more by sample size than by the presence of a meaningful or clinically relevant difference (Sullivan & Feinn, 2012). Alternatively, the effects may be real but modest in magnitude, reflecting subtle and spatially distributed neurophysiological alterations that are difficult to detect at the scalp level. This possibility is particularly relevant in the context of psychiatric disorders, which are highly heterogeneous and may involve overlapping pathophysiological mechanisms (Marquand et al., 2016). Group-level comparisons may therefore obscure meaningful effects present only within specific subgroups or phenotypic dimensions (Marquand et al., 2016). Furthermore, the EEG features selected for analysis may not optimally capture the underlying neural mechanisms relevant to the condition. A mismatch between the features analyzed and the pathophysiological targets of interest could contribute to the weak effects observed. In addition, low signal-to-noise ratio, variability in data quality, and unmodeled sources of variance (e.g., age, sex, comorbidity, medication, and acquisition site) may further reduce sensitivity to true effects. While the persistent pursuit of small effects in underpowered studies has contributed to a vast literature, yet few findings have yielded effect sizes sufficient to support clinical translation (Kapur et al., 2012). Our findings suggest that conventional case-control comparisons may have inherent limitations for EEG biomarker discovery in psychiatry. While larger samples improved the stability of statistical estimates, the resulting effects remained uniformly small. This pattern may reflect the substantial biological heterogeneity present within broad psychiatric diagnoses, whereby meaningful neurophysiological alterations are confined to specific subgroups or symptom dimensions and become diluted in group-average analyses. Consequently, increasing sample size alone may not be sufficient to reveal clinically meaningful biomarkers if the underlying populations remain biologically heterogeneous. Rather than relying on case-control paradigms aimed at diagnosing DSM-defined categories, the field may benefit more from identifying biologically homogeneous subtypes that transcend conventional diagnostic boundaries (Kapur et al., 2012). This shift in focus would prioritize the development of tools that reveal biologically distinct subgroups within broad diagnostic categories, predict differential treatment responses, or clarify disease progression. Methodological frameworks such as unsupervised clustering and normative modeling offer promising opportunities for capturing heterogeneity and identifying clinically relevant subtypes. This realignment in research strategy may ultimately prove more valuable for translational impact than continued efforts to detect marginal effects within ill-defined diagnostic categories.

A further observation from our analysis is the limited diagnostic specificity of EEG alterations. Features showing significant group differences were rarely confined to a single diagnostic category; instead, many appeared across multiple conditions. This aligns with prior findings highlighting the transdiagnostic nature of EEG biomarkers (Newson & Thiagarajan, 2019; Pascucci et al., 2025). Such overlap presents a key barrier to clinical translation, as the utility of a biomarker depends not only on statistical significance but also on its ability to support differential diagnosis, prognosis, or treatment selection. When a feature is shared across several distinct disorders or functional domains, its value within the current categorical nosology is substantially diminished. That said, transdiagnostic markers are not without potential. If they capture shared pathophysiological mechanisms or enable stratification along symptom dimensions rather than diagnostic categories, they may offer meaningful insights, particularly within dimensional frameworks such as those proposed by the Research Domain Criteria (RDoC) initiative (Cuthbert, 2014; Insel et al., 2010). As previously mentioned, rather than replacing existing diagnostic systems, such biomarkers could supplement them by informing individualized treatment strategies and identifying subgroups with distinct neurobiological profiles (Kapur et al., 2012).

It is worth noting that while our study leveraged an unusually large EEG dataset by current standards, cohorts of ∼500 participants per group remain modest compared to the much larger fMRI and MRI cohorts now available. Substantially larger EEG datasets will therefore be needed to achieve comparable power and generalizability. For anxiety and learning disorders, smaller sample sizes further limited generalizability. Furthermore, as sample size approaches the maximum available, subsampling variability naturally decreases due to overlapping samples across iterations, which may slightly bias reproducibility threshold estimates at the upper end of the range.

In summary, our analyses underscore the limitations of underpowered case-control designs in psychiatric EEG research. Small sample sizes lead to unstable and inconsistent group comparisons, undermining the reliability of reported effects. Even with larger samples, observed effect sizes remained uniformly small, raising important questions about the practical utility of conventional group-level approaches for biomarker discovery.

## Data availability

ABCCT and femaleASD datasets can be requested from the NIMH Data Archive platform; MIPDB and HBN datasets are accessible at Child Mind Institute website; LausanneASD dataset can be made available upon request from BR and NC.

## Code availability

Codes are available at https://github.com/MINDIG-1/SNES_CaseControl.git. We used the *pingouin* python package (Vallat, 2018) for statistical calculations, ‘*MNE*’ python package (Gramfort et al., 2013) (https://mne.tools/stable/index.html) for EEG signal processing and custom python-based scripts for the remaining analysis and visualization.

## Acknowledgment

This work was funded by MINDIG as part of its R&D activity. It was also supported by the “Region Bretagne”, Inno R&D project no. 23001155 and Rennes Metropole (AICE project) and the INCR (PsyNorm and Creapark projects). We would like to thank all the researchers who shared their data in open-access and all the participants (patients and controls) who approved the use of their data in research.

## Authors Contribution

A.E., M.H., and S.A. conceived the study and oversaw data analysis and interpretation. A.E. drafted the manuscript and generated the figures with critical revisions from all authors. S.A. assisted with data preprocessing and feature extraction. S.K. initiated the implementation of the project during her internship. B.R. and N.C. provided access to ASD datasets.

## Ethics declaration

### Ethics approval

Ethical oversight was provided by the Chesapeake Institutional Review Board, Yale Institutional Board (Yale, SCRI), and the *Commission Cantonale d’Ethique de la Recherche sur l’être humain* (CER-VD, Switzerland). All procedures performed were in accordance with the ethical standards of the 1964 Helsinki Declaration and its later amendments or comparable ethical standards. All participants (or their legal guardians) provided written informed consent prior to inclusion in the study.

## References

Alexander, L. M., Escalera, J., Ai, L., Andreotti, C., Febre, K., Mangone, A., Vega-Potler, N., Langer, N., Alexander, A., Kovacs, M., Litke, S., O’Hagan, B., Andersen, J., Bronstein, B., Bui, A., Bushey, M., Butler, H., Castagna, V., Camacho, N., … Milham, M. P. (2017). An open resource for transdiagnostic research in pediatric mental health and learning disorders. Scientific Data, 4(1), 170181. 10.1038/sdata.2017.181

Appelhoff, S., Hurst, A. J., Lawrence, A., Li, A., Mantilla Ramos, Y. J., O’Reilly, C., Xiang, L., & Dancker, J. (2022). PyPREP: A Python implementation of the preprocessing pipeline (PREP) for EEG data. (Version 0.4.2) [Computer software]. Zenodo. 10.5281/zenodo.6363576

Arns, M., Conners, C. K., & Kraemer, H. C. (2013). A Decade of EEG Theta/Beta Ratio Research in ADHD: A Meta-Analysis. Journal of Attention Disorders, 17(5), 374–383. 10.1177/1087054712460087

Bigdely-Shamlo, N., Mullen, T., Kothe, C., Su, K.-M., & Robbins, K. A. (2015). The PREP pipeline: Standardized preprocessing for large-scale EEG analysis. Frontiers in Neuroinformatics, 9. 10.3389/fninf.2015.00016

Button, K. S., Ioannidis, J. P. A., Mokrysz, C., Nosek, B. A., Flint, J., Robinson, E. S. J., & Munafò, M. R. (2013). Power failure: Why small sample size undermines the reliability of neuroscience. Nature Reviews Neuroscience, 14(5), 365–376. 10.1038/nrn3475

Clayson, P. E., Carbine, K. A., Baldwin, S. A., & Larson, M. J. (2019). Methodological reporting behavior, sample sizes, and statistical power in studies of event-related potentials: Barriers to reproducibility and replicability. Psychophysiology, 56(11), e13437. 10.1111/psyp.13437

Cohen, J. (2009). Statistical power analysis for the behavioral sciences (2. ed., reprint). Psychology Press.

Cortese, S., Solmi, M., Michelini, G., Bellato, A., Blanner, C., Canozzi, A., Eudave, L., Farhat, L. C., Højlund, M., Köhler-Forsberg, O., Leffa, D. T., Rohde, C., De Pablo, G. S., Vita, G., Wesselhoeft, R., Martin, J., Baumeister, S., Bozhilova, N. S., Carlisi, C. O., … Correll, C. U. (2023). Candidate diagnostic biomarkers for neurodevelopmental disorders in children and adolescents: A systematic review. World Psychiatry, 22(1), 129–149. 10.1002/wps.21037

Cuthbert, B. N. (2014). The RDoC framework: Facilitating transition from ICD/DSM to dimensional approaches that integrate neuroscience and psychopathology: Forum - The Research Domain Criteria Project. World Psychiatry, 13(1), 28–35. 10.1002/wps.20087

Cuthbert, B. N., & Insel, T. R. (2013). Toward the future of psychiatric diagnosis: The seven pillars of RDoC. BMC Medicine, 11(1), 126. 10.1186/1741-7015-11-126

Dede, A. J. O., Xiao, W., Vaci, N., Cohen, M. X., & Milne, E. (2025). Exploring EEG resting state differences in autism: Sparse findings from a large cohort. Molecular Autism, 16(1), 13. 10.1186/s13229-025-00647-3

Ebadi, A., Allouch, S., Mheich, A., Tabbal, J., Kabbara, A., Robert, G., Lefebvre, A., Iftimovici, A., Rodríguez-Herreros, B., Chabane, N., & Hassan, M. (2025). Beyond homogeneity: Charting the landscape of heterogeneity in neurodevelopmental and psychiatric electroencephalography. Translational Psychiatry, 15(1), 223. 10.1038/s41398-025-03441-0

Fortin, J.-P., Cullen, N., Sheline, Y. I., Taylor, W. D., Aselcioglu, I., Cook, P. A., Adams, P., Cooper, C., Fava, M., McGrath, P. J., McInnis, M., Phillips, M. L., Trivedi, M. H., Weissman, M. M., & Shinohara, R. T. (2018). Harmonization of cortical thickness measurements across scanners and sites. NeuroImage, 167, 104–120. 10.1016/j.neuroimage.2017.11.024

Gordillo, D., da Cruz, J. R., Chkonia, E., Lin, W.-H., Favrod, O., Brand, A., Figueiredo, P., Roinishvili, M., & Herzog, M. H. (2023). The EEG multiverse of schizophrenia. Cerebral Cortex, 33(7), 3816–3826. 10.1093/cercor/bhac309

Gramfort, A., Luessi, M., Larson, E., Engemann, D., Strohmeier, D., Brodbeck, C., Goj, R., Jas, M., Brooks, T., Parkkonen, L., & Hämäläinen, M. (2013). MEG and EEG data analysis with MNE-Python. Frontiers in Neuroscience, 7. https://www.frontiersin.org/articles/10.3389/fnins.2013.00267

Insel, T., Cuthbert, B., Garvey, M., Heinssen, R., Pine, D. S., Quinn, K., Sanislow, C., & Wang, P. (2010). Research Domain Criteria (RDoC): Toward a New Classification Framework for Research on Mental Disorders. American Journal of Psychiatry, 167(7), 748–751. 10.1176/appi.ajp.2010.09091379

Jaramillo-Jimenez, A., Tovar-Rios, D. A., Mantilla-Ramos, Y.-J., Ochoa-Gomez, J.-F., Bonanni, L., & Brønnick, K. (2024). ComBat models for harmonization of resting-state EEG features in multisite studies. Clinical Neurophysiology, 167, 241–253. 10.1016/j.clinph.2024.09.019

Jas, M., Engemann, D. A., Bekhti, Y., Raimondo, F., & Gramfort, A. (2017). Autoreject: Automated artifact rejection for MEG and EEG data. NeuroImage, 159, 417–429. 10.1016/j.neuroimage.2017.06.030

Kakuszi, B., Szuromi, B., Tóth, M., Bitter, I., & Czobor, P. (2024). Alterations in resting-state gamma-activity is adults with autism spectrum disorder: A High-Density EEG study. Psychiatry Research, 339, 116040. 10.1016/j.psychres.2024.116040

Kapur, S., Phillips, A. G., & Insel, T. R. (2012). Why has it taken so long for biological psychiatry to develop clinical tests and what to do about it? Molecular Psychiatry, 17(12), 1174–1179. 10.1038/mp.2012.105

Lakens, D. (2013). Calculating and reporting effect sizes to facilitate cumulative science: A practical primer for t-tests and ANOVAs. Frontiers in Psychology, 4. 10.3389/fpsyg.2013.00863

Langer, N., Ho, E. J., Alexander, L. M., Xu, H. Y., Jozanovic, R. K., Henin, S., Petroni, A., Cohen, S., Marcelle, E. T., Parra, L. C., Milham, M. P., & Kelly, S. P. (2017). A resource for assessing information processing in the developing brain using EEG and eye tracking. Scientific Data, 4(1), 170040. 10.1038/sdata.2017.40

Levy, W. J. (1987). Effect of Epoch Length on Power Spectrum Analysis of the EEG. Anesthesiology, 66(4), 489–495. 10.1097/00000542-198704000-00007

Lubba, C. H., Sethi, S. S., Knaute, P., Schultz, S. R., Fulcher, B. D., & Jones, N. S. (2019). catch22: CAnonical Time-series CHaracteristics. Data Mining and Knowledge Discovery, 33(6), 1821–1852. 10.1007/s10618-019-00647-x

Mammarella, I. C., Cardillo, R., & Semrud-Clikeman, M. (2022). Do comorbid symptoms discriminate between autism spectrum disorder, ADHD and nonverbal learning disability? Research in Developmental Disabilities, 126, 104242. 10.1016/j.ridd.2022.104242

Marek, S., Tervo-Clemmens, B., Calabro, F. J., Montez, D. F., Kay, B. P., Hatoum, A. S., Donohue, M. R., Foran, W., Miller, R. L., Hendrickson, T. J., Malone, S. M., Kandala, S., Feczko, E., Miranda-Dominguez, O., Graham, A. M., Earl, E. A., Perrone, A. J., Cordova, M., Doyle, O., … Dosenbach, N. U. F. (2022). Reproducible brain-wide association studies require thousands of individuals. Nature, 603(7902), 654–660. 10.1038/s41586-022-04492-9

Marquand, A. F., Rezek, I., Buitelaar, J., & Beckmann, C. F. (2016). Understanding heterogeneity in clinical cohorts using normative models: Beyond case-control studies. Biological Psychiatry, 80(7), 552–561.

McPartland, J. C., Bernier, R. A., Jeste, S. S., Dawson, G., Nelson, C. A., Chawarska, K., Earl, R., Faja, S., Johnson, S. P., Sikich, L., Brandt, C. A., Dziura, J. D., Rozenblit, L., Hellemann, G., Levin, A. R., Murias, M., Naples, A. J., Platt, M. L., Sabatos-DeVito, M., … Autism Biomarkers Consortium for Clinical Trials. (2020). The Autism Biomarkers Consortium for Clinical Trials (ABC-CT): Scientific Context, Study Design, and Progress Toward Biomarker Qualification. Frontiers in Integrative Neuroscience, 14, 16. 10.3389/fnint.2020.00016

Murray, S. O., Seczon, D. L., Pettet, M., Rea, H. M., Woodard, K. M., Kolodny, T., & Webb, S. J. (2025). Increased alpha power in autistic adults: Relation to sensory behaviors and cortical volume. Autism Research, 18(1), 56–69. 10.1002/aur.3266

Neo, W. S., Foti, D., Keehn, B., & Kelleher, B. (2023). Resting-state EEG power differences in autism spectrum disorder: A systematic review and meta-analysis. Translational Psychiatry, 13(1), 389. 10.1038/s41398-023-02681-2

Newson, J. J., & Thiagarajan, T. C. (2019). EEG Frequency Bands in Psychiatric Disorders: A Review of Resting State Studies. Frontiers in Human Neuroscience, 12, 521. 10.3389/fnhum.2018.00521

Pascucci, D., Menétrey, M. Q., Passarotto, E., Luo, J., Paramento, M., & Rubega, M. (2025). EEG brain waves and alpha rhythms: Past, current and future direction. Neuroscience & Biobehavioral Reviews, 176, 106288. 10.1016/j.neubiorev.2025.106288

Pelphrey, K. (2012). Multimodal Developmental Neurogenetics of Females with ASD. NIMH Data Repositories. 10.15154/4DAT-5683

Pion-Tonachini, L., Kreutz-Delgado, K., & Makeig, S. (2019). ICLabel: An automated electroencephalographic independent component classifier, dataset, and website. NeuroImage, 198, 181–197. 10.1016/j.neuroimage.2019.05.026

Poil, S.-S., Bollmann, S., Ghisleni, C., O’Gorman, R. L., Klaver, P., Ball, J., Eich-Höchli, D., Brandeis, D., & Michels, L. (2014). Age dependent electroencephalographic changes in attention-deficit/hyperactivity disorder (ADHD). Clinical Neurophysiology, 125(8), 1626–1638. 10.1016/j.clinph.2013.12.118

Precenzano, F., Parisi, L., Lanzara, V., Vetri, L., Operto, F. F., Pastorino, G. M. G., Ruberto, M., Messina, G., Risoleo, M. C., Santoro, C., Bitetti, I., & Marotta, R. (2020). Electroencephalographic Abnormalities in Autism Spectrum Disorder: Characteristics and Therapeutic Implications. Medicina, 56(9), 419. 10.3390/medicina56090419

Reddan, M. C., Lindquist, M. A., & Wager, T. D. (2017). Effect Size Estimation in Neuroimaging. JAMA Psychiatry, 74(3), 207. 10.1001/jamapsychiatry.2016.3356

Richardson, J. T. E. (2011). Eta squared and partial eta squared as measures of effect size in educational research. Educational Research Review, 6(2), 135–147. 10.1016/j.edurev.2010.12.001

Rodríguez-Herreros, B., Mheich, A., Osório, J. M. A., Richetin, S., Junod, V., Arnold, L., Aeschbach, V., Adamou, L. N., Mendes, L., Gschwend, K., Romascano, D., Yu, P., Hassan, M., & Chabane, N. (2025). Social, cognitive and sensory dimensions of cortical network overconnectivity in young children with autism spectrum disorder (p. 2025.02.12.637813). bioRxiv. 10.1101/2025.02.12.637813

Sullivan, G. M., & Feinn, R. (2012). Using Effect Size—Or Why the *P* Value Is Not Enough. Journal of Graduate Medical Education, 4(3), 279–282. 10.4300/JGME-D-12-00156.1

Tabbal, J., Ebadi, A., Mheich, A., Kabbara, A., Güntekin, B., Yener, G., Paban, V., Gschwandtner, U., Fuhr, P., Verin, M., Babiloni, C., Allouch, S., & Hassan, M. (2025). Characterizing the heterogeneity of neurodegenerative diseases through EEG normative modeling. Npj Parkinson’s Disease, 11(1), 1–12. 10.1038/s41531-025-00957-6

Tabbal, J., Ebadi, A., Robert, G., Rodríguez-Herreros, B., Mheich, A., Chabane, N., Lefebvre, A., Iftimovici, A., Allouch, S., & Hassan, M. (2025). Transdiagnostic electrophysiological subtypes reveal brain-behavior dimensions in youth psychiatry. Neuroscience. 10.1101/2025.08.01.668189

Vallat, R. (2018). Pingouin: Statistics in Python. Journal of Open Source Software, 3(31), 1026. 10.21105/joss.01026

van Diessen, E., Numan, T., van Dellen, E., van der Kooi, A. W., Boersma, M., Hofman, D., van Lutterveld, R., van Dijk, B. W., van Straaten, E. C. W., Hillebrand, A., & Stam, C. J. (2015). Opportunities and methodological challenges in EEG and MEG resting state functional brain network research. Clinical Neurophysiology: Official Journal of the International Federation of Clinical Neurophysiology, 126(8), 1468–1481. 10.1016/j.clinph.2014.11.018

Yun, S. (2024). Advances, challenges, and prospects of electroencephalography-based biomarkers for psychiatric disorders: A narrative review. Journal of Yeungnam Medical Science, 41(4), 261–268. 10.12701/jyms.2024.00668

