## Supplementary Materials for "Sample size critically shapes the reliability of EEG case-control findings in psychiatry"

### Methods

#### Dataset

**Table. S1 | Mean age (± standard deviation) by group.**

| Group | HC | ADHD | ASD | ANX | LD |
|---|---|---|---|---|---|
| **Mean age ± SD** | 10.16±3.23 | 9.82±2.93 | 9.84±2.99 | 10.88±2.91 | 9.85±2.68 |

**Table S2 | Distribution of subjects across diagnostic groups and datasets, including sex proportions within each group–dataset combination**

| Group | HBN | ABCCT | MIPDB | femaleASD | LausanneASD | Total |
|---|---|---|---|---|---|---|
| **HC** | 251 (46% M) | 118 (69% M) | 59 (61% M) | 101 (53% M) | 29 (38% M) | 558 (54% M) |
| **ADHD** | 1213 (27% M) | - | 8 (75% M) | - | - | 1221 (28% M) |
| **ASD** | 173 (14% M) | 268 (77% M) | - | 101 (52% M) | 12 (83% M) | 554 (53% M) |
| **ANX** | 305 (52% M) | - | - | - | - | 305 (52% M) |
| **LD** | 236 (46% M) | - | - | - | - | 236 (46% M) |
| **Total** | 2178 (34% M) | 386 (75% M) | 67 (63% M) | 202 (53% M) | 41 (51% M) | **2874** (42% M) |

**Table. S3 | Number of subjects with multiple diagnoses across the groups considered in the study**

| Primary Diagnosis | ADHD | ADHD +ANX | ADHD +ANX+ LD | ADHD +LD | ANX | ANX +LD | ASD | ASD+ ADHD | ASD+ ADHD +ANX | ASD+ADHD+ANX+LD | ASD+ ADHD +LD | ASD+ ANX | ASD+ ANX+ LD | ASD+ LD | LD |
|---|---|---|---|---|---|---|---|---|---|---|---|---|---|---|---|
| **ADHD** | 677 | 182 | 60 | 202 | 0 | 0 | 0 | 107 | 31 | 10 | 24 | 0 | 0 | 0 | 0 |
| **ANX** | 0 | 74 | 26 | 0 | 156 | 27 | 0 | 0 | 9 | 4 | 0 | 10 | 2 | 0 | 0 |
| **ASD** | 0 | 0 | 0 | 0 | 0 | 0 | 48 | 74 | 29 | 9 | 15 | 6 | 2 | 7 | 0 |
| **LD** | 0 | 0 | 27 | 27 | 0 | 29 | 0 | 0 | 0 | 3 | 0 | 0 | 0 | 3 | 147 |

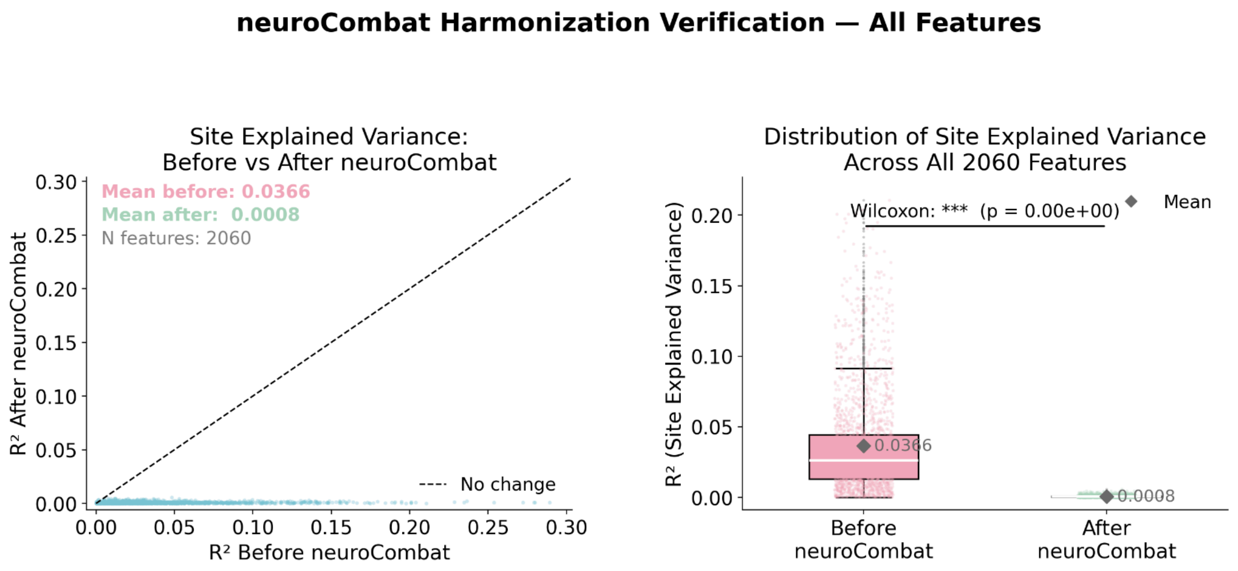


**Fig. S1 | Verification of neuroCombat harmonization across all 2,060 EEG feature columns. Left column:** scatter plots of explained variance before versus after neuroCombat for site (top, $\eta^2$). Each point represents one feature. **Right column**: boxplots showing the distribution of explained variance before and after neuroCombat; diamond markers indicate the mean. Site-explained variance was reduced from a mean $\eta^2$ of 0.037 to 0.001 (~98% reduction), confirming effective removal of site-related variance.

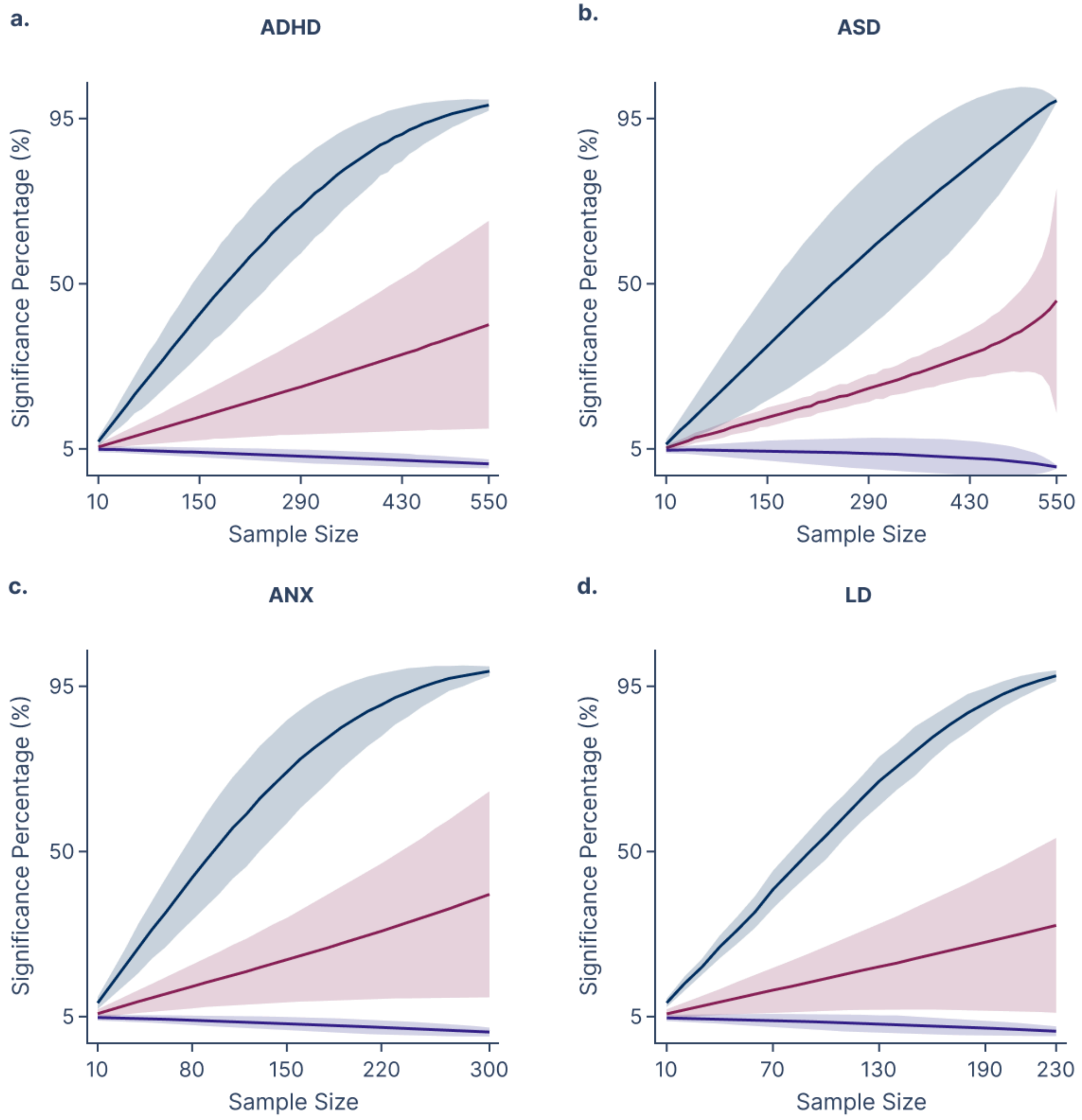


**Fig. S2 | Average significance percentage for all features across diagnostic groups as a function of sample size.** Each panel shows the mean significance percentage (proportion of 1,000 resampling iterations yielding $p < 0.05$) averaged across all features, plotted as a function of subsample size for each diagnostic group: a, ADHD, b, ASD, c, ANX, and d, LD. The dark blue line represents features that reached the 95% consistency threshold at the full sample size (truly significant features), the purple line represents features that remained below the 5% consistency threshold across all sample sizes, and the red line represents features that did not conform to either of these conditions. Shaded areas indicate the standard deviation across features within each category.

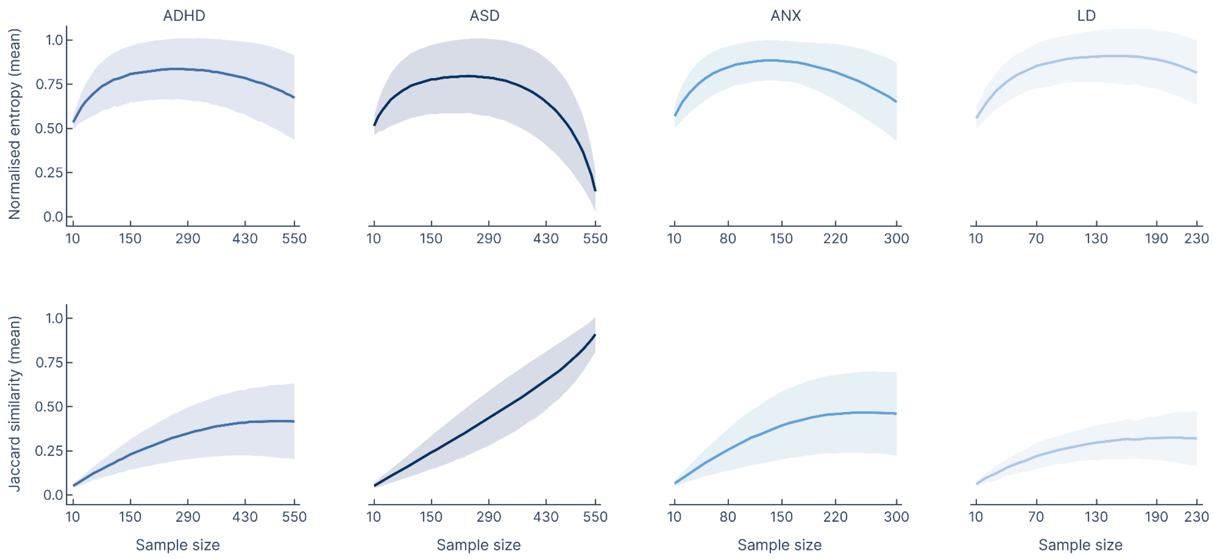


**Fig. S3 | Spatial reproducibility metrics as a function of sample size across diagnostic groups. Top row:** mean normalised entropy of significant feature counts, averaged across features and computed over 1,000 subsampling iterations, plotted against sample size for each diagnostic group (ADHD, ASD, ANX, LD). Higher entropy indicates greater variability in which channels are identified as significant across iterations. **Bottom row:** mean Jaccard similarity between pairs of significant feature sets, averaged across features, reflecting the consistency of feature identification across iterations. In both rows, shaded regions represent the standard deviation. Across the 103 feature families, 36, 65, 33, and 12 showed at least one significant channel for ADHD, ASD, ANX, and LD, respectively.

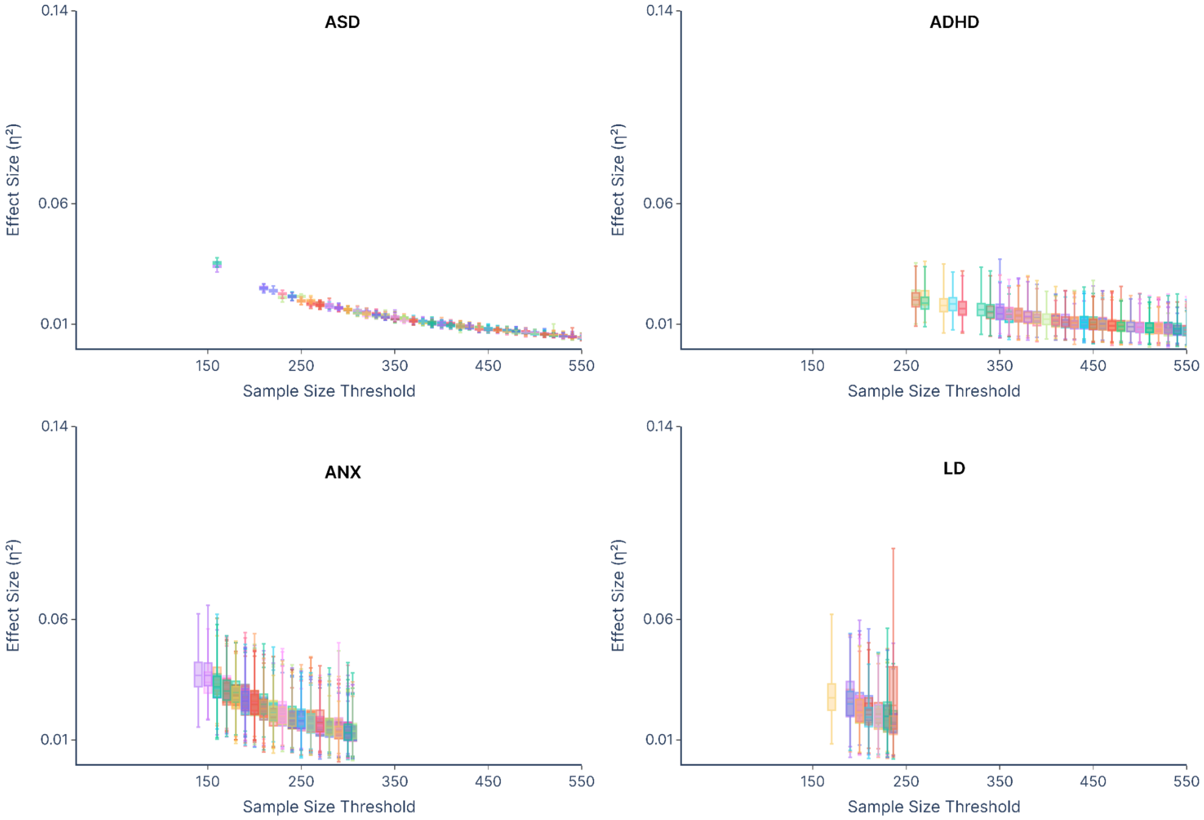


**Figure S4 | Distribution of full-sample effect sizes ($\eta^2$) plotted against sample size thresholds for all four diagnostic groups (ASD, ADHD, ANX, LD).** Each box represents the distribution of effect sizes across 1,000 subsampling iterations for a single significant feature at a given threshold; all significant features for each group are shown.
